# Characterization of scar tissue biomechanics during adult murine flexor tendon healing

**DOI:** 10.1101/2021.11.09.467960

**Authors:** Antonion Korcari, Mark R. Buckley, Alayna E. Loiselle

**Affiliations:** Center for Musculoskeletal Research, University of Rochester Medical Center, Rochester, NY 14642; Department of Biomedical Engineering, University of Rochester, Rochester, NY

**Keywords:** Biomechanics, tendon injury, regenerative healing, scar tissue, function

## Abstract

Tendon injuries are very common and result in significant impairments in mobility and quality of life. During healing, tendons produce a scar at the injury site, characterized by abundant and disorganized extracellular matrix and by permanent deficits in mechanical integrity compared to healthy tendon. Although a significant amount of work has been done to understand the healing process of tendons and to develop potential therapeutics for tendon regeneration, there is still a significant gap in terms of assessing the direct effects of therapeutics on the functional and material quality specifically of the scar tissue, and thus, on the overall tendon healing process. In this study, we focused on characterizing the mechanical properties of only the scar tissue in flexor digitorum longus (FDL) tendons during the proliferative and remodeling healing phases and comparing these properties with the mechanical properties of the composite healing tissue. Our method was sensitive enough to identify significant differences in structural and material properties between the scar and tendon-scar composite tissues. To account for possible inaccuracies due to the small aspect ratio of scar tissue, we also applied inverse finite element analysis (iFEA) to compute mechanical properties based on simulated tests with accurate specimen geometries and boundary conditions. We found that the scar tissue linear tangent moduli calculated from iFEA were not significantly different from those calculated experimentally at all healing timepoints, validating our experimental findings, and suggesting the assumptions in our experimental calculations were accurate. Taken together, this study first demonstrates that due to the presence of uninjured stubs, testing composite healing tendons without isolating the scar tissue overestimates the material properties of the scar itself. Second, our scar isolation method promises to enable more direct assessment of how different treatment regimens (e.g., cellular ablation, biomechanical and/or biochemical stimuli, tissue engineered scaffolds) affect scar tissue function and material quality in multiple different types of tendons.

## Introduction

Tendons are highly organized fibrous musculoskeletal tissues that function by transmitting muscle-generated forces to the bones and enabling skeletal movement. Tendon injuries are very common and account for more than 30% of all musculoskeletal consultations^1,2^. In the US alone, there are more than 300,000 tendon and ligament repair surgeries performed annually^3^. After injury, tendons demonstrate a scar-mediated reparative response rather than a regenerative healing phenotype and produce fibrotic tissue around the healing region that is characterized by abundant and disorganized extracellular matrix (ECM). The persistent fibrotic tissue is mechanically inferior compared to the native tendon, and results in permanent impairment of full tendon function^6–8^ due to a combination of insufficient restoration of mechanical properties, which significantly increases the risk of re-injury^9^, and the formation of peritendinous adhesions or connections between the tendon and surrounding tissues which impairs tendon movement^10,11^.

A tremendous amount of work has focused on improving our understanding of the biology of tendon homeostasis and healing, and on identifying potential regenerative approaches to improve tendon healing. For example, many studies have focused on the role of different tendon ECM components (e.g., collagen V, VI, decorin biglycan) in the maintenance of tendon homeostasis and during healing^12–17^. Other studies have focused on improving tendon healing by utilizing different biochemical molecules or signaling pathways such as the NF-kB pathway, TGF-β, aspirin, and Tenomodulin^4,10,18–24^. Finally, an increasing number of studies have tried to understand and exploit the functions of different tendon and non-tendon cell populations (resident tendon cells, adipose-derived stem cells, induced pluripotent stem cells, immune cells) to achieve tendon regeneration during healing^4,21,25–29^. Unfortunately, although the above studies have offered a significant amount of multidisciplinary knowledge and may identify future potential therapeutic approaches, there is still no consensus therapy in clinical use. Therefore, pre-clinical animal models continue to have great importance with respect to better defining the molecular and cellular mediators of the healing process, and in understanding how different therapeutics modulate healing, with an important emphasis on understanding how specific treatment modalities alter the tendon mechanical environment.

Uniaxial tensile testing is the “gold” standard method to assess the physiologically relevant function and material quality of healing tendons. Traditionally, at the post-injury timepoint of interest the tendon tissue is dissected, cleaned by removing any extra non-tendon tissues, gripped by the 2 tendon edges, placed in some type of buffer e.g., 1X Phosphate Buffer Solution (PBS) to keep the tendon hydrated, and is uniaxially tensile stretched in a displacement-controlled manner until failure. Typically, the tendon gauge length involves a composite tissue of both un-injured tendon adjacent to the injury/repair site and the healing scar tissue region where the injury occurred^18,30–37^. However, up to this point, the potential relative mechanical contribution of only the scar tissue/healing region during tendon healing is unknown. Moreover, given that the scar tissue represents the mechanically weakest region in the healing tendon^38,39^, a more precise characterization of the scar tissue structural and material properties will likely provide important insights in the efficacy of a treatment regimen in terms of improved tendon function and material quality.

In the present study, our first objective was to successfully characterize the structural (CSA, peak load, stiffness) and material (peak stress, tangent modulus) properties specifically of the tendon scar tissue during adult FDL tendon healing in a sensitive and robust way. Next, to validate the assumptions of our experimental calculations, we more directly computed material properties using inverse finite element modeling (iFEM) with accurate specimen geometries and boundary conditions. We hypothesized that the scar tissue of FDL tendons will exhibit different structural and material properties compared to the tendon-scar composite tissue at multiple healing timepoints.

## Materials and Methods

### Mice

All studies were approved by the University Committee on Animal Resources. Wildtype mice on a C57Bl/6J background were used for this study. All mouse studies were performed with 10-12-week-old male and female mice. All mouse work (surgeries, harvests) was performed in the morning. Mice were kept on a 12-hour light/dark cycle.

### Flexor Tendon Repair

Complete transection and repair of the murine flexor digitorum longus (FDL) tendon was performed as previously described^40^. Briefly, mice were injected prior to surgery with 15-20μg of sustained-release buprenorphine analgesic. Mice were anesthetized with Ketamine (100mg/kg) and Xylazine (4mg/kg). Following sterilization of the surgery region, the flexor tendon was transected at the myotendinous junction (MTJ) to reduce early-stage strain-induced rupture of the repair site and the skin was closed with a 5-0 suture. A small incision was made on the posterior surface of the hindpaw, the FDL tendon was located and completely transected using micro spring-scissors. The tendon was repaired using 8-0 suture and the skin was closed with a 5-0 suture. Following surgery, animals resumed prior cage activity, food intake, and water consumption.

### Scar and tendon-scar composite tissue dissection and preparation for mechanical testing

The FDL tendon was carefully separated from the calf muscle under a dissecting microscope. The tarsal tunnel was then cut and the FDL tendon was slowly released from the tarsal tunnel and isolated until just proximal to the healing region. Next, the distal FDL was cut and separated at the bifurcation of the digits and isolated proximally until just distal to the healing region. Finally, the healing region was carefully isolated, and the intact healing FDL tendon was placed in a new petri dish. Under the dissecting microscope, any additional connective tissues were removed, and the tendon-scar composite FDL tendon was prepared for testing. Scar tissue was evident in all dissected FDL tendons.

To isolate only the scar tissue and grip it for uniaxial tensile testing, two pieces of sandpaper were placed on each end of the scar tissue and glued together using cyanoacrylate (Superglue, LOCTITE) to grip it in such a way where the gauge length of the gripped tissue included only the scar and not any ‘native’ tendon proximal or distal to the repair site. For gripping of the tendon-scar composite tissue, approximately 1.3-1.5 mm of native tendon was included proximal and distal to the repair site. All the steps above were performed with the tissue being constantly submerged in PBS to avoid any potential tissue drying.

### Quantification of scar and tendon-scar composite tissue dimensions

Each gripped sample was transferred into a semi-customized uniaxial microtester (eXpert 4000 MicroTester, ADMET, Inc., Norwood MA). The microtester, along with the sample, was transferred to an inverted microscope (Olympus BX51, Olympus) to visualize the tendon and quantify the gauge length, the width, and the thickness. The gauge length of each sample was set as the end-to-end distance between the sandpapers. The cross-section of the tendon was assumed to be an ellipse, where the width of the tissue represents the major axis, while the thickness of the tissue represents the minor axis of the ellipse, as previously described^41,42^. To quantify the width and thickness of the scar and the tendon-scar composite groups, three sub-equal measurements for each parameter were taken across the scar tissue (proximal, central, and distal scar) and across the tendon-scar composite tissue (proximal tendon, scar, and distal tendon), and the average value of the width and thickness were set as the representative values, respectively.

### Uniaxial tensile testing

A uniaxial displacement-controlled stretching of 1% strain per second until failure was applied for both the scar tissue and the tendon-scar composite tissue by utilizing the uniaxial microtester after quantifying the dimensions of each tested tendon (gauge length, CSA). Load and grip-grip displacement data were recorded, and the failure mode was tracked for each mechanically tested sample. All samples from both groups (scar and tendon-scar composite tissues) at D14 and D21 timepoints failed at the midsubstance after the uniaxial tensile stretch. The tensile stress was taken to equal the recorded load divided by the initial CSA, while the tensile strain was taken to equal the recorded grip-grip displacement divided by the gauge length. The load-displacement and stress-strain data were plotted and analyzed to determine structural (*stiffness, peak load*) and material (*modulus, peak stress*) properties. The slope of the linear region from the load displacement graph during the tensile testing was determined to be the stiffness of the tested sample. The maximum load recorded from the load-displacement graph during the tensile testing was determined to be the peak load parameter of the tested sample. The slope of the linear region from the stress-strain graph during the tensile testing was taken to equal the tangent modulus parameter of each tested tendon. Note that this calculation assumes that stress and strain are uniform within each specimen. Finally, the maximum stress value recorded from the stress-strain graph during the tensile testing was determined to be the peak stress (or strength) of the tested tendon tissue. The structural properties provide information about the mechanical integrity of the tendon which depends on the tendon cross-sectional area. The material properties provide information about the intrinsic material quality of the tissue independently of the cross-sectional area, and thus can be used as a metric outcome of tendon function.

### Finite Element Model Construction and iFE analysis for linear tangent modulus quantification

Because the tested specimens had small aspect ratios (gauge length/width ~ 1.1 for D14 and 1.7 for D21), stress and strain distributions induced during tensile testing were presumably nonuniform (according to Saint-Venant’s principle). Therefore, it was necessary to test the validity of our assumption that modulus can be computed from the slope of the overall tissue stress versus tissue strain curve in the linear region. To this end, 3D finite element models of each experimentally tested FDL scar tissue at both D14 and D21 post-surgery were created **[Figure 4. B, C]**. First, elliptic cylinders with accurate dimensions (gauge length, width, and thickness) matching each tested specimen were generated in OnShape, Next, each scar geometry was imported into FEBio and meshed. The number of elements in each model was 14,084 and the number of nodes was 3,205. To accurately reflect the effects of specimen gripping, fixed boundary conditions (in all directions) were applied at one end of the scar, while the moving end was fixed perpendicular to the direction of stretching (i.e., perpendicular to the long axis of the elliptic cylinder).

Each experimental mechanical test was simulated in FEBio starting from the beginning of the linear region. To identify the exact start and end point of the linear region, we applied a linear regression model to our experimental data, quantified the corresponding R^2^ value, and only groups of points with values of R^2^ ≥ 0.995 were accepted as part of the linear region **[Figure 4. A]**. The recorded displacement (relative to the beginning of the linear region) was applied to moving end of the elliptic cylinder in the finite element model.

The scar tissue was modeled as a neo-Hookean material. Because we restricted our simulations to the linear region of the stress-strain curve, we chose a constitutive relationship model that has linear behavior for small strains. Because scar tissue is poorly aligned (**Fig. 1A, B**), we chose an isotropic model.

**Figure 1.**
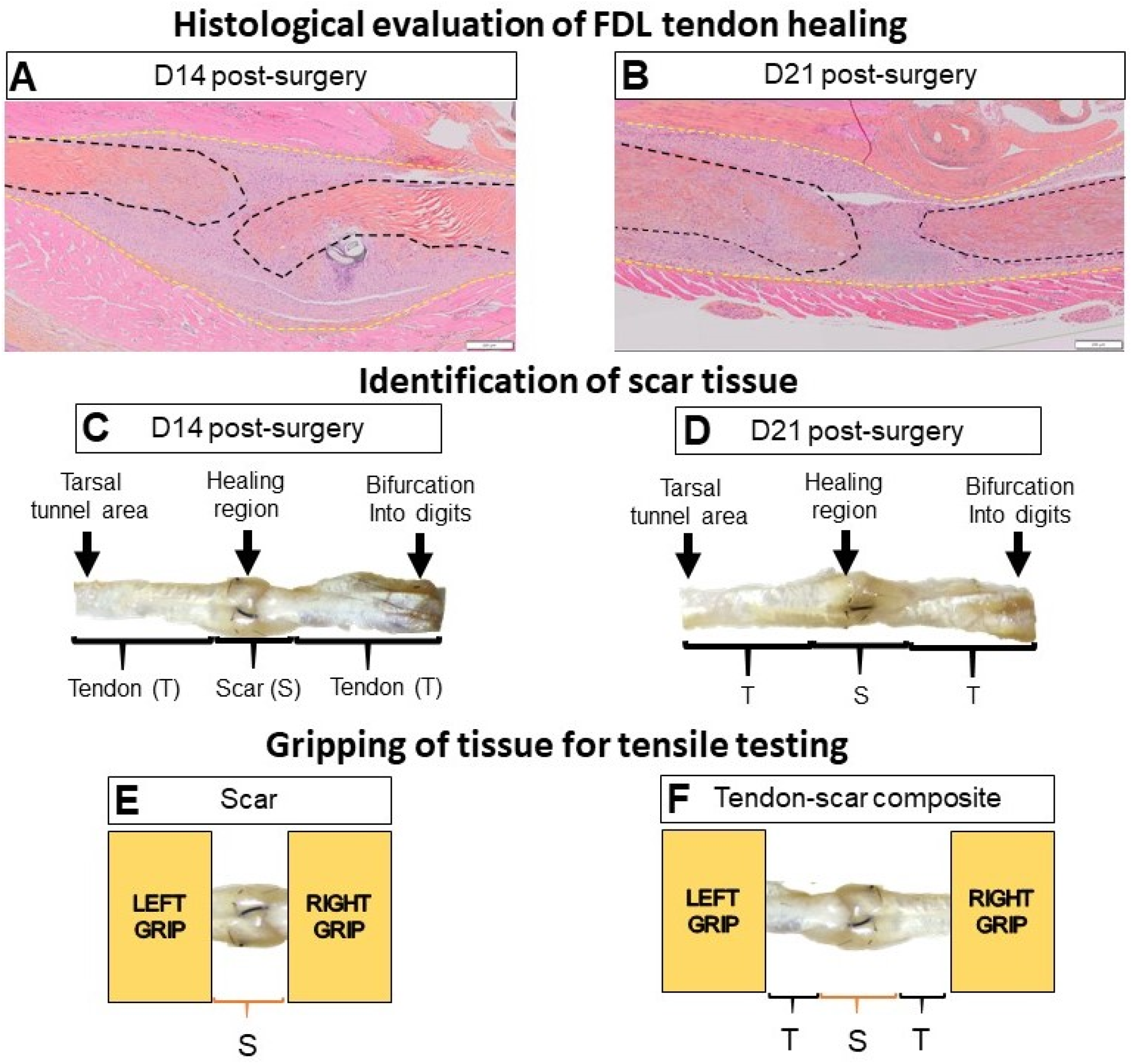
FDL tendon scar and tendon-scar composite tissues can be successfully identified, gripped, and prepared for uniaxial tensile testing. **A**. ABHOG staining on FDL tendons at D14 post-surgery. **B**. ABHOG staining on FDL tendons at D21 post-surgery. **C**. A representative harvested healing tendon at D14 post-surgery. **D**. A representative harvested healing tendon at D21 post-surgery. The scar tissue can be easily identified approximately in the middle of the tissue (healing region). The increase of the width and thickness in the healing region due to the scar formation is evident. The anatomical markers used to harvest each tissue are the region where the tendon bifurcates into digits and the region where the tendon goes through the tarsal tunnel area. After dissection under the microscope, **(E)** only the scar tissue or **(F)** a composite of tendon-scar-tendon tissue was gripped on each end using sandpaper and superglue and prepared for uniaxial tensile testing.

To calculate the tangent modulus from the simulations, we utilized an iterative optimization approach (inverse finite element analysis, or iFEA) implemented in FEBio. Specifically, the modulus of the simulated scar was assumed as an initial arbitrary number, the above stretching simulation was performed, and the simulated reaction force at the fixed boundary was quantified. The correct/true value of the modulus for each simulation was identified when the simulated reaction force from FEBio matched (difference between FEBio-simulated and experimental reaction force <1%) the experimental force at the end of the linear region. Next, we recorded the coordinates (force, displacement) in both the start and end of each linear region which represented the specific load and displacement each scar tissue could withstand before entering the plastic deformation.

### Histological evaluation of tendon healing

Hindpaws containing the repaired FDL tendon were harvested at D14 and D21 post-repair (n=4 per time-point) as previously described^29,43^. Briefly, the hindpaw was disarticulated at the distal tibia, skin on the top of the foot was removed, and samples were fixed in 10% neutral buffered formalin for 72h with the ankle fixed at ~90°. The tissue was then decalcified for 14 days and processed for paraffin histology. Five-micron serial sections were cut through sagittal plane of the foot and stained with Alcian Blue/ Hematoxylin/ Orange G (ABHOG). Stained slides were digitally imaged using a VS120 Virtual Slide Microscope (Olympus).

### Statistical analysis

Quantitative data was analyzed via GraphPad Prism and is presented as mean ± standard deviation (SD). A student’s t-test was used to analyze the gauge length and mechanical properties data between the scar and the tendon-scar composite tissue., as well as between the experimentally calculated and FEBio-based tangent moduli. A one-way ANOVA followed by Tukey’s post-hoc analysis was used to analyze the width and thickness data between the three different regions in the scar and the tendon-scar composite tissue. p values ≤ 0.05 were considered significant. * indicates p<0.05, ** indicates p<0.01, *** indicates p<0.001, **** indicates p<0.0001.

## Results

### Scar tissue is shorter and more homogeneous in width and thickness compared to the composite tissue

Morphological staining of the healing tendons was consistent with our previous characterization of the FDL healing process^4,20,21,40,44^, and demonstrated the formation of bridging scar tissue between the ends of the native tendon at the repair site on D14 and D21 (**Figure 1A &B**), suggesting potential partial restoration of mechanical intergrity. Morphologically, the thickness of the repair site is decreased from D14 to D21 and ECM alignment is enhanced, consistent with tissue remodeling, and increased integration of the healing tissue-native tendon is observed. Consistent with our histology, we dissected the healing tendon scar tissue along with extra proximal and distal tendon. Dissected tendons were gripped using sandpaper on both ends, their dimensions were quantified, and tensile testing was performed.

To quantify the dimensions of the scar and the tendon-scar composite groups, as described above, three sub-equal measurements were taken across the tissues and the average value of the width and thickness were set as their representative values, respectively **(Figure 2A &B)**. At D14 post-surgery, the scar tissue had a 63% shorter gauge length compared to the tendon-scar composite tissue (p<0.0001) **(Figure 2C)**. The scar tissue had a relatively homogeneous width (proximal scar =1.88 ± 0.049mm; central scar =1.9 ±0.046mm; distal scar =1.88 ± 0.05mm) and thickness (proximal scar =1.51 ± 0.139mm; central scar =1.54 ± 0.141mm; distal scar =1.52 ± 0.141mm) **(Figure 2D &E)**. In contrast, the tendon-scar composite tissue had a more heterogeneous width with proximal tendon (1.03 ± 0.097mm) being significantly lower compared to scar tissue (1.7 ± 0.029mm, p<0.0001) and distal tendon (1.53 ± 0.042mm, p<0.001), while scar tissue was significantly higher than distal tendon (p<0.01) **(Figure 2F)**. Similarly, the thickness was also heterogeneous, with proximal tendon (0.95 ± 0.088mm) being significantly lower compared to scar tissue (1.52 ± 0.029mm, p<0.0001) and distal tendon (1.26 ± 0.044mm, p<0.001), while scar tissue was significantly higher than distal tendon (p<0.01) **(Figure 2G)**.

**Figure 2.**
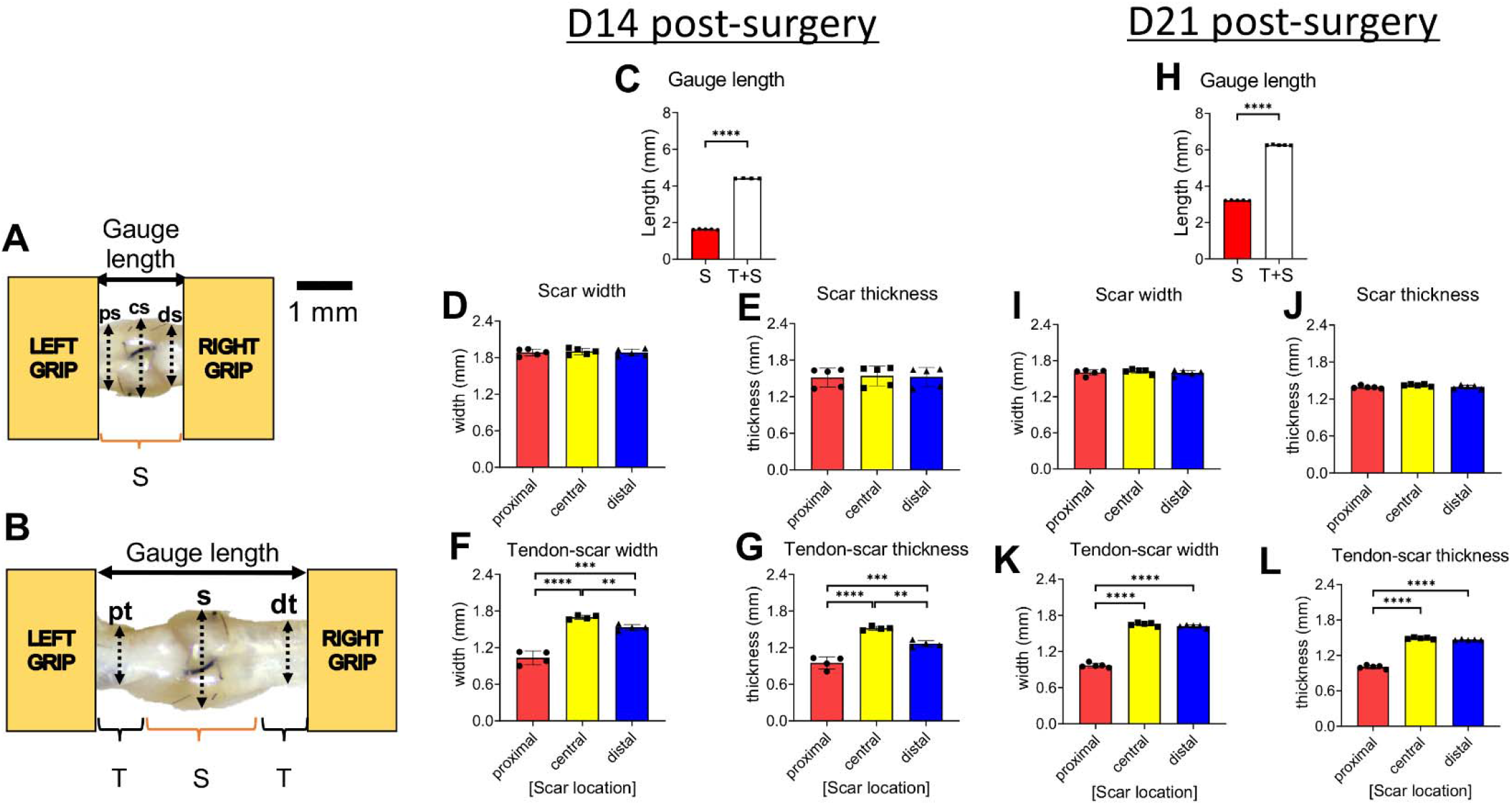
Quantification of dimensions (gauge length, width, thickness) of the scar and tendon-scar composite tissues prior to mechanical testing. **A**. The gauge length of the scar tissue and the three sub-equal regions, proximal scar (ps), central scar (cs), and distal scar (ds), used to quantify the width and the thickness of the scar tissue. **B**. The gauge length of the tendon-scar composite tissue and the three sub-equal regions, proximal tendon (pt), scar (s), distal tendon (dt), used to quantify the width and the thickness of the composite tissue. **C**. Gauge length between the scar and tendon-scar composite tissue at D14 post-surgery. **D**. The width of the scar tissue at D14 post-surgery. **E**. The thickness of the scar tissue at D14 post-surgery. **F**. The width of the tendon-scar composite tissue at D14 post-surgery. **G**. The thickness of the tendon-scar composite tissue at D14 post-surgery. **H**. Gauge length between the scar and tendon-scar composite tissue at D21 post-surgery **I**. The width of the scar tissue at D21 post-surgery **J**. The thickness of the scar tissue at D21 post-surgery. **K**. The width of the tendon-scar composite tissue at D21 post-surgery. **L**. The thickness of the tendon-scar composite tissue at D21 post-surgery. N=4-5 per group. Statistical significance between regions was determined using a one-way ANOVA followed by a Tukey’s post-hoc analysis. ** indicative of p < 0.01; *** indicative of p < 0.001; **** indicative of p < 0.0001

At 21 days post-surgery, the scar tisue was approximately 49% shorter than the tendon-scar composite tissue (p<0.00001) **(Figure 2H)**. Both the width (proximal scar =1.6 ±0.043mm; central scar =1.62 ± 0.034mm; distal scar =1.6 ± 0.038mm) and thickness (proximal scar =1.39 ± 0.02mm; central scar =1.42 ± 0.0149mm; distal scar =1.39 ± 0.026mm) **(Figure 2I &J)** of the scar tissue were homogeneous, consistent with 14 days post-surgery. In contrast, the tendon-scar composite tissue exhibited a more heterogeneous width with proximal tendon (0.97 ± 0.034mm) being significantly lower compared to scar tissue (1.65 ± 0.025mm) (p<0.0001) and distal tendon (1.62 ± 0.021mm, p<0.0001) **(Figure 2K)**. Similarly, the thickness of the tendon-scar composite tissue was heterogeneous with proximal tendon (1 ± 0.027mm) being significantly lower compared to scar tissue (1.49 ± 0.014mm, p<0.0001) and distal tendon (1.47 ± 0.008mm, p<0.0001) **(Figure 2L)**.

### Scar tissue has significantly different CSA and linear tangent moduli compared to tendon-scar composite tissue at both D14 and D21 post-surgery

At D14 post-surgery, the scar tissue CSA was approximately 35% larger (p<0.01) compared to the tendon-scar composite tissue due to the heterogeneity in dimensions of the different regions in the tendon-scar composite tissue compared to the more homogeneous scar tissue **(Figure 3A)**. The peak load and stiffness between the scar and the tendon-scar composite tissues showed no significant differences (p>0.05) **(Figures 3B &C)**. The peak stress values between the scar and the tendon-scar composite tissues did not show any significant differences (p>0.05) **(Figure 3D)**, while the tangent modulus of the scar tissue was almost 6-fold less than that of the tendon-scar composite tissue (p<0.01) **(Figure 3E)**.

**Figure 3.**
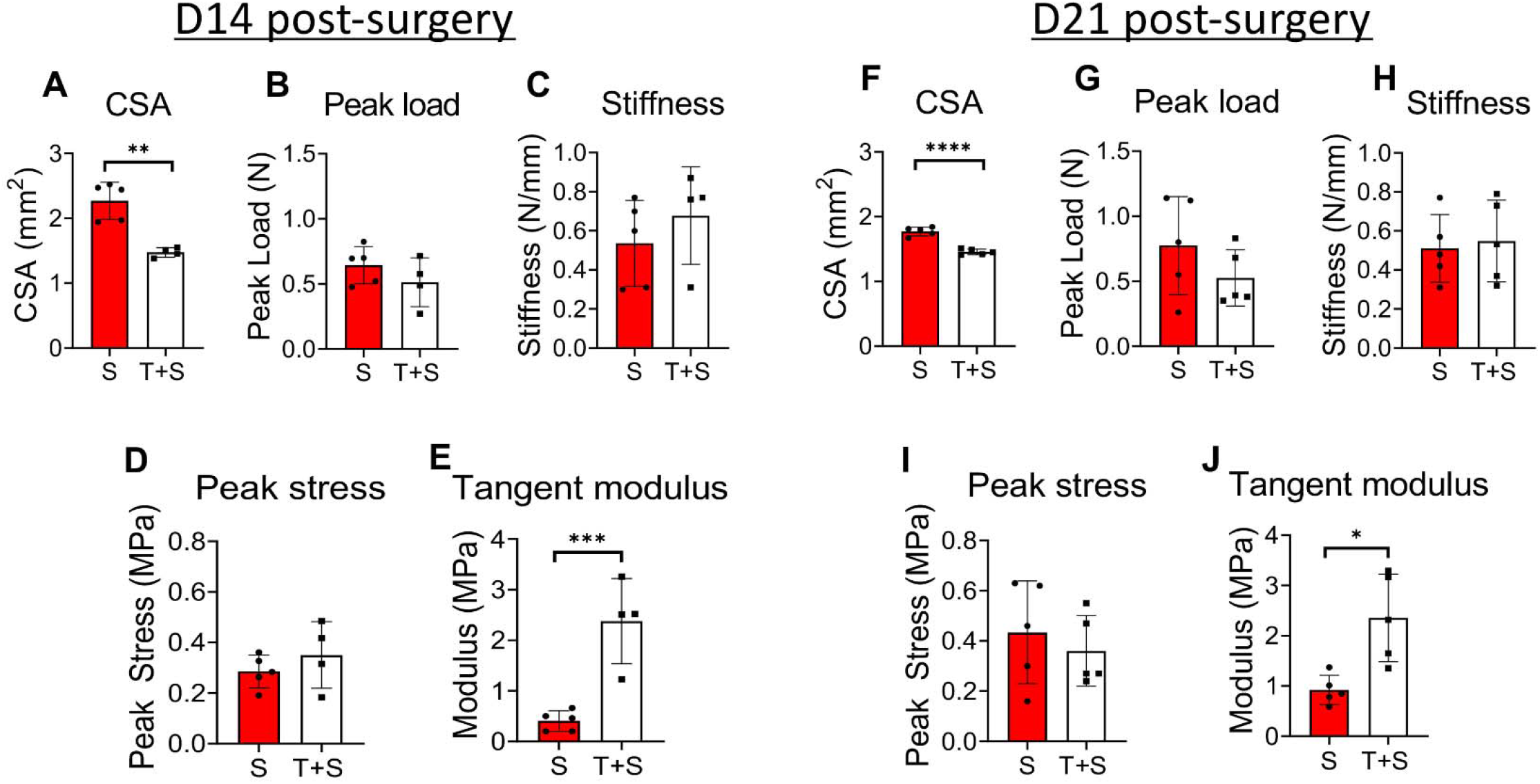
The scar tissue has significantly different cross-sectional area (CSA) and material quality compared to the tendon-scar composite tissue at both 14 and 21 days post-surgery. **A**. The CSA of the scar (S) tissue is significantly higher compared to the tendon-scar composite (T+S) tissue at 14 days post-surgery. **B**. The peak load between the S and T+S tissues is similar at 14 days post-surgery. **C**. The stiffness between the S and T+S tissues is similar at 14 days post-surgery. **D**. The peak stress between the S and the T+S tissues is similar at 14 days post-surgery. **E**. The tangent modulus of the S tissue is significantly lower compared to the T+S tissue at 14 days post-surgery. **F**. The CSA of the S tissue is significantly higher compared to the T+S tissue at 21 days post-surgery. **G**. The peak load between the S and T+S tissues is similar at 21 days post-surgery. **H**. The stiffness between the S and T+S tissues is similar at 21 days post-surgery. N=4-5 per group. Statistical significance between groups was determined using Student’s t-test. **I**. The peak stress between the scar S and the T+S tissues is similar at 21 days post-surgery. **J**. The tangent modulus of the S tissue is significantly lower compared to the T+S tissue at 21 days post-surgery. N=4-5 per group. Statistical significance between regions was determined using unpaired Student t-test. * indicative of p < 0.05; *** indicative of p < 0.001.

Similarly, at D21 post-surgery, the scar tissue CSA was 17.77% larger (p<0.00001) compared to the tendon-scar composite tissue **(Figure 3F)**, while both the peak load stiffness between the scar and the tendon-scar composite tissues did not show any significant differences (p>0.05) **(Figure 3G &H)**. The peak stress between the scar and the tendon-scar composite tissues did not show any significant differences (p>0.05) **(Figure 3I)**, while the tangent modulus of the scar tissue was almost 2.6-fold less than that of the tendon-scar composite tissue (p<0.01) **(Figure 3J)**.

### Validation of experimentally calculated scar tissue linear tangent moduli values via FEBio

At D14 post-surgery, the tangent modulus of the scar tissue calculated experimentally was slightly lower than the tangent modulus calculated using iFEA. However, this difference was not statistically significant (p=0.2046) **(Figure 4D)**. In a similar manner, at D21 post-surgery, FEA-based tangent modulus values were slightly higher than the experimentally calculated tangent modulus values. However, this difference was also not statistically significant (p=0.3383) **(Figure 4E)** and was less pronounced than in the D14 group.

**Figure 4.**
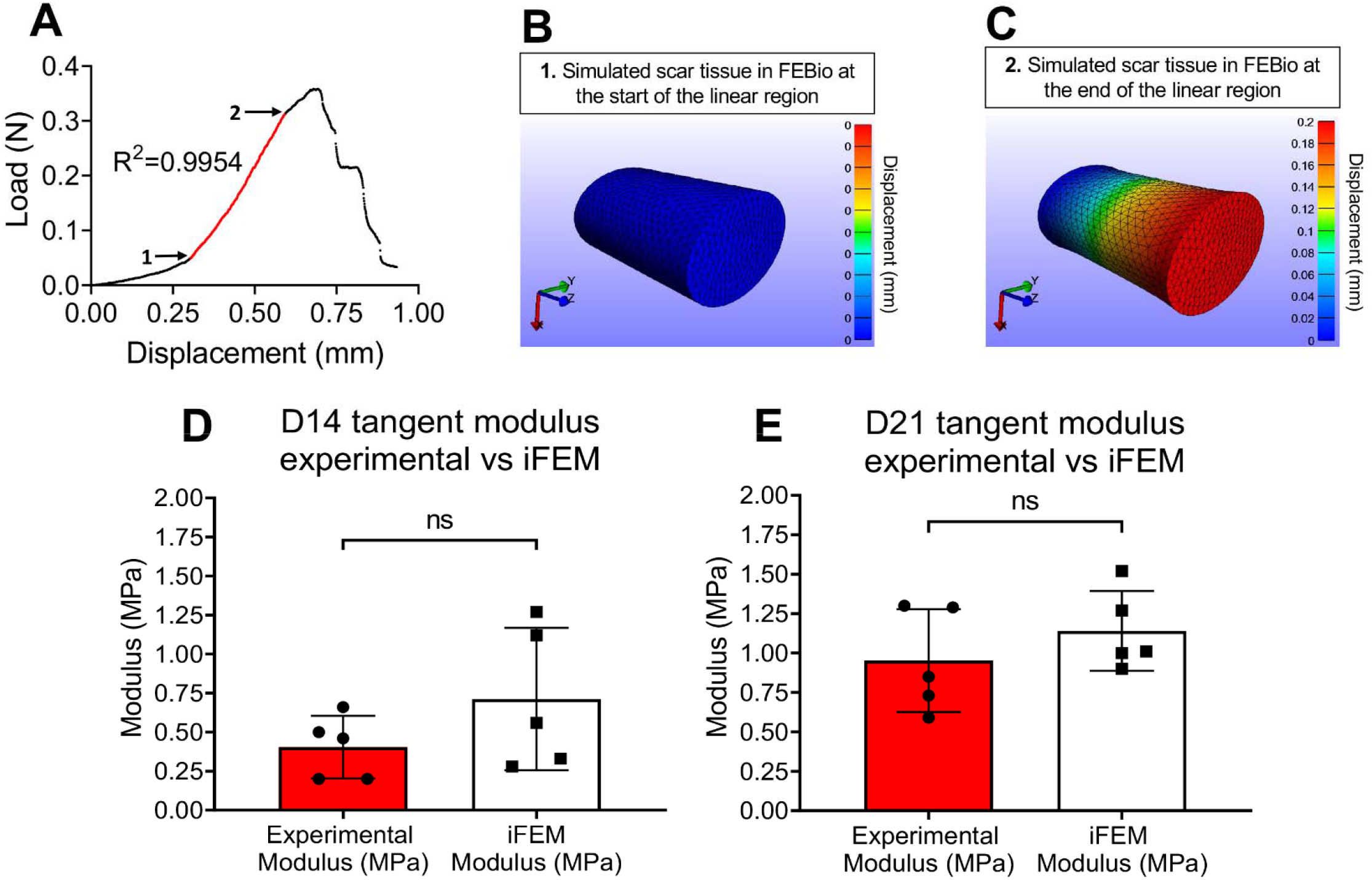
Validation of experimentally calculated scar tissue tangent moduli using a Finite Elements Analysis (FEA) modeling approach. **A**. Representative load-displacement graph of a scar tissue where the linear region was identified, values of force and displacement for the start and endpoints of the linear region were recorded and used to simulate the linear region of a displacement-controlled uniaxial tensile stretch in FEBio. **B**. A representative simulated scar tissue in FEBio right at the beginning of the linear region. **C**. A representative simulated scar tissue in FEBio after the simulated uniaxial stretching right at the end of the linear region. **D**. The tangent moduli calculated using the experimental data (Experimental Modulus) versus FEBio (iFEM Modulus) at D14 post-surgery were found not to be significantly different from each other. **E**. The tangent moduli calculated using the experimental data (Experimental Modulus) versus FEBio (iFEM Modulus) at D21 post-surgery were found not to be significantly different from each other. N=4-5 per group. Statistical significance between regions was determined using unpaired Student t-test. ns indicative of not statistically significant.

**Supplemental figure 1.**
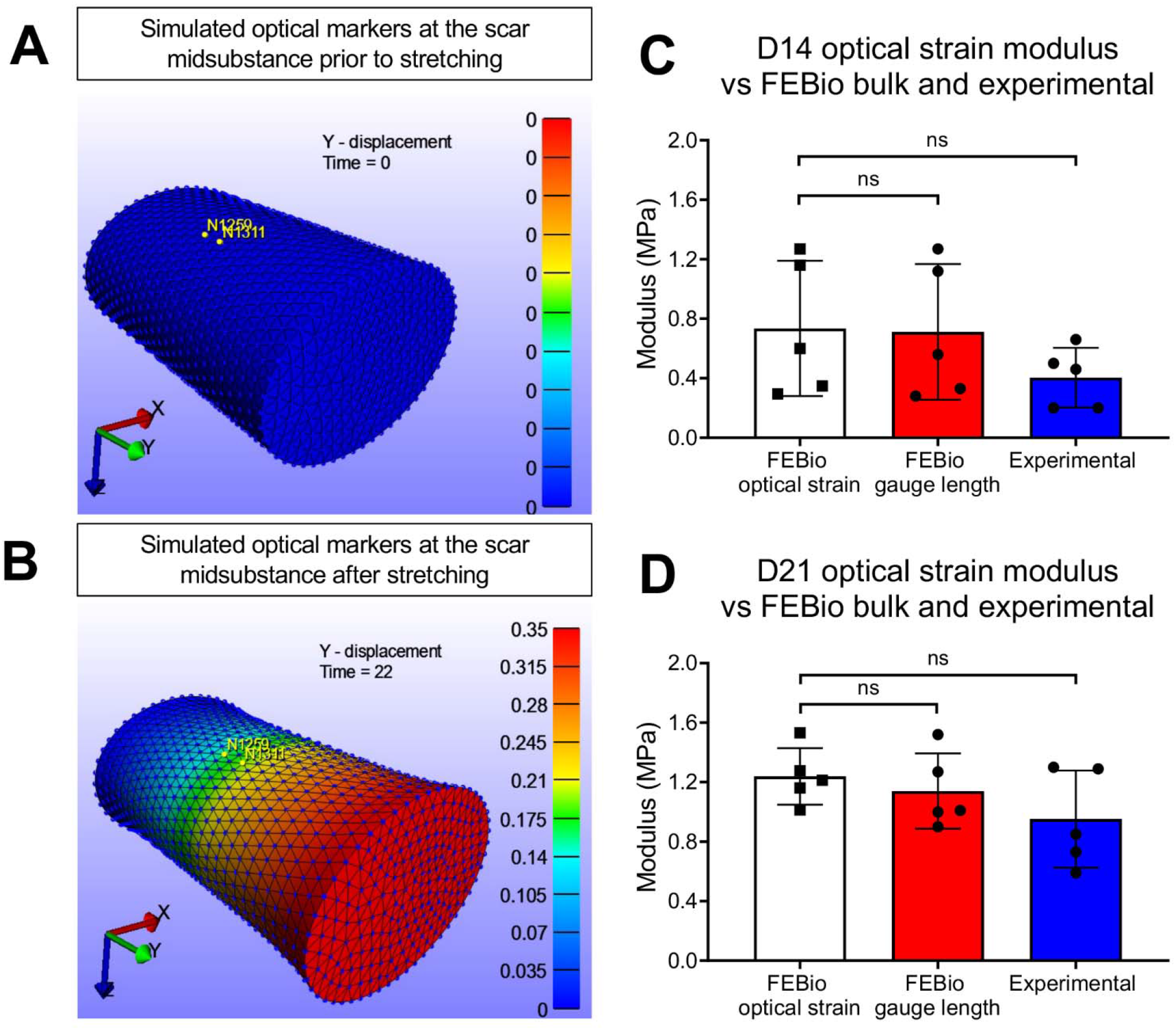
Validation of both experimental and FEBio gauge length-based linear tangent moduli via optical strain markers-based tangent moduli quantification. **A**. FEBio-based simulated scar tissue with optical markers at the midsubstance of the sample prior to tensile stretching. **B**. FEBio-based simulated scar tissue with optical markers at the midsubstance of the sample right after the tensile stretching. **C**. Comparison of the tangent moduli calculated via FEBio-based optical strain markers to tangent moduli calculated via FEBio-based gauge length and experimentally at D14 post-surgery. **D**. Comparison of the tangent moduli calculated via FEBio-based optical strain markers to tangent moduli calculated via FEBio-based gauge length and experimentally at D21 post-surgery.

## Discussion

In this study, our objective was to successfully characterize the mechanical properties of the isolated scar tissue at different timepoints of healing. Next, we utilized a finite elements approach using the FEBio program to simulate the uniaxial tensile stretching of scar tissues and validated the results we generated experimentally in terms of tissue material quality. We were able to show the importance of dissecting the intrinsic structural and material properties of only the scar tissue and to underscore that with such a method, it will be possible to ‘screen’ different treatment regimens (e.g. cellular ablation, biomechanical and/or biochemical stimuli, tissue engineered scaffolds) and assess whether a specific treatment results in a more regenerative or fibrotic response of the healing tendon by directly quantifying the structural and material properties of the scar tissue. Such mechanical characterization is important because it gives us insight on the direct effect of a treatment regimen on the scar tissue biomechanics. In contrast, mechanical characterization of a composite tendon-scar tissue will result in an inaccurate understanding of the mechanical environment of the scar tissue since it is the mechanically weakest part of the tendon^38,39^ and when combined with native non-injured tissue, the sensitivity for detecting any mechanical changes in the scar is drastically minimized. We found that the scar tissue exhibits a significant increase in CSA and a significant decrease in tangent modulus compared to the tendon-scar composite at both 14- and 21-days post-surgery. To our knowledge, this is the first study to successfully quantify the mechanical properties of tendon scar tissue and to demonstrate significant differences between scar and tendon-scar composite tissue structural and material properties. These findings highlight the importance of choosing a representative tendon gauge length for mechanical assessment during healing. Characterization of the mechanical properties of only the scar tissue, whenever possible, is a direct assessment of the function and material quality of the healing region and might be a better indicator of a more regenerative vs fibrotic healing response of the tendon.

At D14 post-surgery (proliferative healing phase), we found that the CSA of the scar tissue group was significantly higher compared to the tendon-scar composite group and similarly the histological evaluation verified such an increase in the scar CSA. At this phase, various intrinsic and extrinsic cell populations are recruited in the healing region and initiate the production of a provisional scar tissue, mainly collagen III and different proteoglycans to close the wound area and provide an initial partial restoration of the tendon mechanical integrity^6,27,45^. Consistent with our histological and quantitative characterization of the scar CSA, the provisional scar tissue is characterized by a significant increase in the CSA and a random deposition and orientation of the different secreted ECM molecules in contrast to the highly organized morphology of the collagen fibers and ECM molecules in the healthy tendon tissue^6,27,45^. Consistent with this, Zhao et al., performed total transection and repair of the flexor digitorum profundus (FDP) tendons in a canine model and they showed that the FDP tendons had a significant increase in the cross-section of the healing region^12,18,33,46^. We also found that at the same time there were no significant differences in both the peak load and stiffness between the two groups. These data indicate the “attempt” of the injured tendon to bear higher amounts of load. This attempt, although it offers a subsequent amount of strength, it has a counter-effect on the function/material quality of the tissue^47–50^ as we demonstrated here by the significant differences of the scar tissue versus the tendon-scar composite tissue tangent moduli at both 14- and 21-days post-surgery.

At D21 post-surgery (early remodeling phase), we found that the CSA of the scar tissue group was significantly higher compared to the tendon-scar composite group. However, the difference of the CSA between the scar and tendon-scar composite tissue at D21 (early remodeling phase) post-surgery was smaller compared to that at D14 (proliferative phase) post-surgery. Also, the scar tissue CSA at D21 post-surgery was smaller than that at D14 post-surgery. These results were further validated by our histological characterization which showed that as we move into the earlier remodeling healing phase, the CSA of the scar seems to decrease. As healing moves into the earlier remodeling phase (D21) or consolidation phase, there is a significant decrease in both the cellularity as well as the production of ECM^6,45^. The collagen III in the healing region starts being replaced by the much more organized collagen I resulting in a slight decrease of the cross-section and in the production of a more fibrous scar tissue^45,50^. In terms of material quality, the scar tissue tangent modulus was significantly lower compared to that of the tendon-scar composite tissue tangent modulus, however, the difference of the tangent modulus between the two groups at D21 post-surgery was smaller compared to that at D14 post-surgery. These data and our histological characterization indicate that as the scar tissue remodeling continues, the newly produced collagen I fibers start to organize better compared to the collagen III across the longitudinal axis of the tendon tissue, resulting in improvements of the function and material quality of the healing tendon^6,45^.

In the healing FDL scar tissue, the aspect ratio (gauge length divided by width) was approximately 1:1 for D14 and ~1.7:1 for D21 post-surgery. Such small aspect ratios could potentially introduce complex stress and strain distributions that impair the accuracy of the uniform stress/strain assumption. To test whether such artifacts were introduced in our mechanically tested scars, we used finite element analysis to simulate our mechanical tests with boundary conditions reflecting the effects of gripping and accurate specimen geometries. This allowed us to use iterative optimization to quantify the “real” tangent modulus (material quality) of the scar independently of artifacts resulting from inaccurate assumptions. By comparing the experimentally calculated versus iFEA-based tangent moduli for scar tissue at D14 and D21, we found no significant differences at all timepoints. Thus, our experimentally calculated moduli reflected the “true” material quality of the scar tissue within the variability of the tested specimens. Of course, gripping-induced spatial variations in stress and strain were certainly present in our experiments and are evident in our simulations (Supplemental Figure 1). However, the magnitude of their effect on the calculated material properties was not substantial compared with the inherent variability of the acquired data (most likely due to biological variations). Interestingly, when we compared tangent moduli calculated experimentally with tangent moduli calculated using IFEA for D14 scar tissue, although significant changes were not observed, the average modulus appeared higher in the iFEA group. However, the relative difference in tangent modulus between experimental and iFEA-based values of the tangent modulus in the D21 group was smaller, likely because the aspect ratio was higher in this group and stress was therefore more uniform.

This study was not without limitations. The model used in our iFE analysis assumed that the tested scar tissue was isotropic and did not account for fiber alignment, collagen uncrimping or collagen realignment. However, we restricted our analysis to the linear region (after collagen uncrimping and realignment have taken place) and only tested scar tissue, which is poorly aligned. In addition, our strain measurements used grip-grip strain instead of optical strain. However, we imaged all specimens on a reflected light microscope before and after testing and did not observe any signs of slippage. Moreover, we simulated our mechanical tests using iFEA with accurate specimen geometries and boundary conditions to confirm the assumptions of our experimental calculations. We also performed FEA to simulate optical strain measurement in our models and found that calculating strain based on tracking marked locations within the tissue would not have yielded statistically distinct values of tangent modulus compared with our approach (Supplemental Figure 1). A further limitation is that we did not look at timepoints beyond D21 of healing, however, the objectives of this study were to show that we can successfully isolate, grip, tensile test, and detect differences in the mechanical properties of only the scar tissue during the different healing phases. We have previously shown that the healing FDL tendons exhibit different healing characteristics between D14- and D21 post-surgery^4,21,43^, indicative of the different healing phases. Time-points prior to D14 were not analyzed due to sample fragility, low mechanical properties and insufficient scar formation. In addition, we have not examined the effects of age or sex on the isolated mechanical properties of scar tissue. Finally, we believe that it might be technically challenging to apply this method to tendons that are short (patellar tendon) or that have a more complicated geometry (supraspinatus tendons) due to the difficulty of isolating and gripping only the scar tissue.

Collectively, these data demonstrate that it is possible to characterize the mechanical properties of only the scar tissue in FDL tendons at multiple healing timepoints. This method was highly sensitive, could identify mechanical differences between scar and tendon-scar composite tissue, and was successfully validated by comparison with iFEA based on models with accurate specimen geometry and grip-simulating boundary conditions. This approach will open new technical avenues in the tendon field by making it possible to more directly assess the efficacy of numerous treatment regimens (e.g., cellular ablation, biomechanical and/or biochemical stimuli, tissue engineered scaffolds) on tendon function and material quality.

## Acknowledgements

This work was supported in part by NIH/ NIAMS R01AR073169 and R21 AR073961 (to AEL), and R01 AR070765 (to MRB). The HBMI and BBMTI Cores were supported by NIH/ NIAMS P30AR069655. The content is solely the responsibility of the authors and does not necessarily represent the official views of the National Institutes of Health.

